# A novel mutation alters the stability of PapA2 resulting in the complete abrogation of sulfolipids in clinical mycobacterial strains

**DOI:** 10.1101/487124

**Authors:** Vipul Panchal, Nidhi Jatana, Anchal Malik, Bhupesh Taneja, Ravi Pal, Apoorva Bhatt, Gurdyal S Besra, Lipi Thukral, Sarika Chaudhary, Vivek Rao

## Abstract

The analysis of whole genome has revealed specific geographical distribution of *Mycobacterium tuberculosis* (Mtb) strains across the globe suggestive of unique host dependent adaptive mechanisms in Mtb evolved out of selective pressures. We provide an important correlation of a genome-based mutation to a molecular phenotype across two predominant clinical Mtb lineages of the Indian subcontinent. We have identified a distinct lineage specific mutation-G247C, translating into an alanine to proline conversion in the *pap*A2 gene of Indo oceanic lineage 1 [L1] Mtb strains; restoration of cell wall sulfolipids by simple genetic complementation of *papA*2 from lineage 3 [L3] or from H37Rv (Lineage 4-L4) attributed the loss of this glycolipid to this specific mutation in Indo oceanic L1 Mtb. Investigation into the structure of Mtb PapA2 revealed a distinct non-ribosomal peptide synthetase (NRPS) C domain conformation with an unconventional presence of Zinc binding motif. Surprisingly, the A83P mutation did not map to either the catalytic centre in the N-terminal subdomain or any of the substrate binding region of the protein. On the contrary, the inherent ability of mutant PapA2 to form insoluble aggregates and molecular simulations with the Wt/ mut PapA2 purports an important role for the surface associated 83^rd^ residue in protein conformation. The present study demonstrates the importance of a critical structural residue in the papA2 protein of Mtb and helps establish a link between observed genomic alteration and its molecular consequence in the successful human pathogen Mtb.

## INTRODUCTION

*Mycobacterium tuberculosis* (Mtb) has the dubious distinction of being one of the most successful human pathogens by virtue of its extreme adaptability and survivability in the face of stress. Being an intracellular pathogen, Mtb has evolved to sense and manipulate the host to its advantage. Whole genome sequencing methods have classified Mtb strains across the globe into 7 major lineages (L1-L7) that have co-evolved with its specific host population and environment (1).

The Mtb cell wall is now recognized as a complex entity unique in its composition of complex polyketide lipids like TDM, SL, DAT-PAT, PDIM. This assembly requires a finely tuned array of metabolic functions involving the biosynthesis, maturation, transport and assembly of precursors from the cytoplasm to the exterior (2–5). The ability of Mtb strains to alter their cell wall repertoire to effectively communicate with host cells, modulate immune signalling and play a pivotal role in intracellular fitness is well recognised (6–10). Unique lipids like PGL in Mtb strains are associated with down-regulation of the inflammatory response and consequent hyper-virulence of the strains (11, 12, 13). Interestingly, minor modifications like cylopropanation of mycolic acids leads to marked alterations in the host immune activation/ suppression (14, 15). Mtb has evolved to manipulate the expression of its lipids as a counter for intracellular stress (16, 17).

Sulfolipids represent Mtb specific lipids that have been the focus of research over the last several years (18–20). The presence of sulfolipids has been classically associated with virulence of mycobacteria (21–24). Moreover, recent evidence has further corroborated their role in bacterial physiology. The biosynthetic pathway of mature sulfolipids in Mtb has been well characterized with the synthesis involving step wise addition of 4 fatty acyl chains to sulfated trehalose by acyl transferases- PapA2 (Rv3820c), PapA1 (Rv3824c) and Chp1(Rv3822) coupled to export of the lipid to the outer cell wall by the transporters-Mmpl8 and Sap (3, 25, 26).

In this study, we have employed whole-genome based analysis to pin point the molecular basis of loss of mature sulfolipid expression in the cell wall of the Indo oceanic Mtb lineage (a subset of Mtb lineage 1. We demonstrate that a G247C SNP in the *papA*2 gene, encoding the first acyl transferase in sulfolipid biosynthesis, results in a detrimental modification of the alanine-83 to proline. Expression of *papA*2 from H_37_Rv or N24 (lineage 3) was sufficient to restore mature sulfolipid in the cell walls of deficient strains establishing that this mutation is solely responsible for the loss of sulfolipid in these strains. By using X ray- crystallography, we demonstrate that PapA2 attains a classical NRPS condensation (C) domain architecture with two subdomains arranged in V shape, each with coenzyme-A dependent acyltranferase (CAT) fold, 2 cross-over points and catalytic centre at the interface of two subdomains. The presence of a distinctive Zn finger motif in the N-terminal region of Mtb PapA2 represents a unique modification of this protein from other known acyl transferases. By MD simulation studies of mutant PapA2, we demonstrate that the A83P mutation induces significant misfolding of the protein resulting in global changes in protein conformation. Coupled to our inability to acquire soluble mutant protein from *E. coli*, we provide evidence for an important role for the surface associated mutation in structural stability of Mtb PapA2 and SL-1 biosynthesis.

## Results

### Genome sequence analysis provides crucial insight into the loss of mature sulfolipid in lipid scaffold of Lineage 1 Mtb strains

Previous studies (27, 28) have implicated a loss of sulfatides from Mtb strains belonging to South India- (Indo oceanic L1). In order to test if the differences also panned to the other predominant strain of the subcontinent – (L3) in Northern India, we investigated the total cell wall associated lipid content among these two Mtb lineages. Fig.1 shows the 2D TLC lipid profiles of 3 strains each from Mtb lineages 1 and 3. A uniform absence of SL-1 from the apolar lipid fraction was the most distinct feature in all the 3 strains of Indo oceanic L1 Mtb (Fig.1A). Most of the other apolar or polar lipids were consistent in both the lineages (Fig. 1B-i-v).

**Fig. 1.**
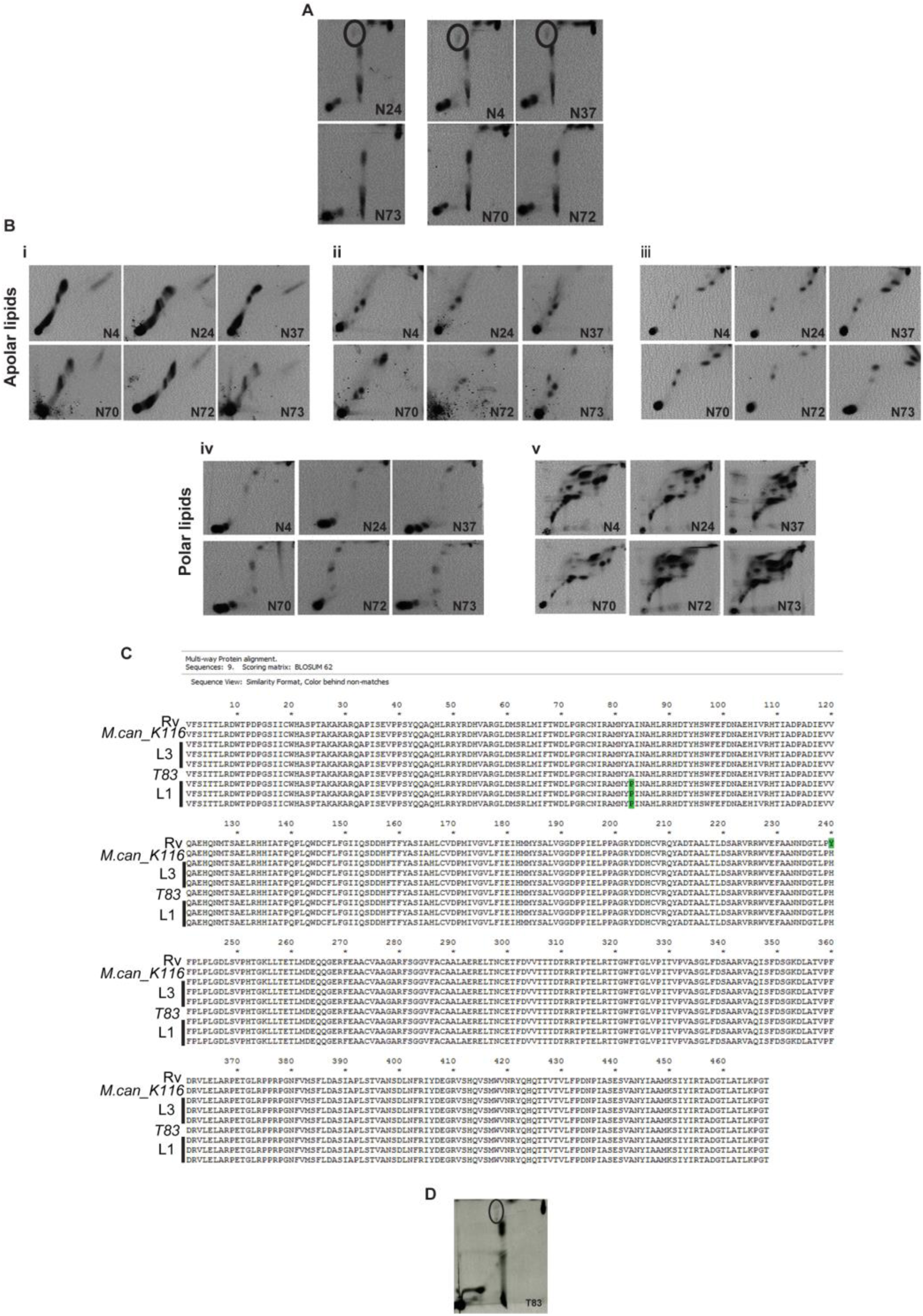
A non-synonymous SNP in PapA2 gene of Indo oceanic Mtb strains is responsible for loss of mature SL-1. (A, B)-analysis of radiolabelled lipids by 2-D TLC: A) Apolar lipids of ^14^C-acetate labelled Mtb Indo oceanic L1 (N73, N70, N72) and L3 (N24, N4, N37) were resolved in solvent D and visualized by radio-imaging. The mature sulfolipid-SL-1 spot is encircled. B) Analysis of total lipids from ^14^C acetate labelled cultures by 2-D TLC in the different solvent systems (i-solvent A, ii-solvent B and iii-solvent C, iv-solvent D and v-solvent E). C) Comparison of PapA2 sequence of the reference Mtb strain (H_37_Rv), *M. canetti K116*, the three strains each of Indo oceanic L1 (N73, N70, N72) and L3 (N24, N4, N37) and the L1 strain from Vietnam – T83. The mutated residue A83P is marked in green. D) Analysis of apolar lipids from Mtb T83 from ^14^C-acetate labelled logarithmic cultures resolved in solvent D is shown. The mature sulfolipid-SL-1 spot is encircled.

In an attempt to understand the molecular basis of this sulfolipid loss in (Indo oceanic-L1), we resorted to genome sequence comparison with previously reported L3 strains and the reference strain H_37_Rv (29). Closer examination of the SNP list pointed towards a common mutation in the Indo oceanic-L1 Mtb genomes (pos. 428579); this G to C conversion resulted in conversion of the 83^rd^ alanine of PapA2 (a polyketide associated acyl transferase involved in sulfolipid biosynthesis of Mtb) to proline (Fig. 1C); a non-tolerable mutation to protein function and structure (SIFT analysis). Interestingly, this mutation was not observed in *M. canetti* or the other closely related L1 strain - T83 (belonging to the Vietnam region) indicating significant specificity of this mutation to the Indian subcontinent (Fig. 1C); consequently, T83 strain was capable of producing mature SL-1 in the cell wall associated non-polar lipid fraction (Fig. 1D).

### Trans-complementation of *papA*2 from L3 genome restores SL-1 biosynthesis in representative L1 strain

The presence of a deleterious non-synonymous mutation, A83P, only in PapA2 allowed us to hypothesise its association with the absence of SL-1 in Indo oceanic lineage-1. In order to test this hypothesis, we ectopically expressed HA-tagged *papA*2 gene from H_37_Rv or L3 in the Indo oceanic L1 strain-N73 and confirmed expression by tag specific immunoblotting (Fig. 2A). We checked for the restoration of SL-1 biosynthesis by 1D as well as 2D -radiometric TLC. Restoration of SL-1 was observed only when *papA*2 was expressed from H_37_Rv or L3, but not when *papA*2 of Indo oceanic L1 or in case of vector control were used (Fig. 2B). Similar restoration of SL-1 in another Indo oceanic L1 strain – N70 explicitly confirms the causative role of A83P mutation in the loss of PapA2 function and consequent SL-1 in this subset of L1 Mtb strains.

**Fig. 2.**
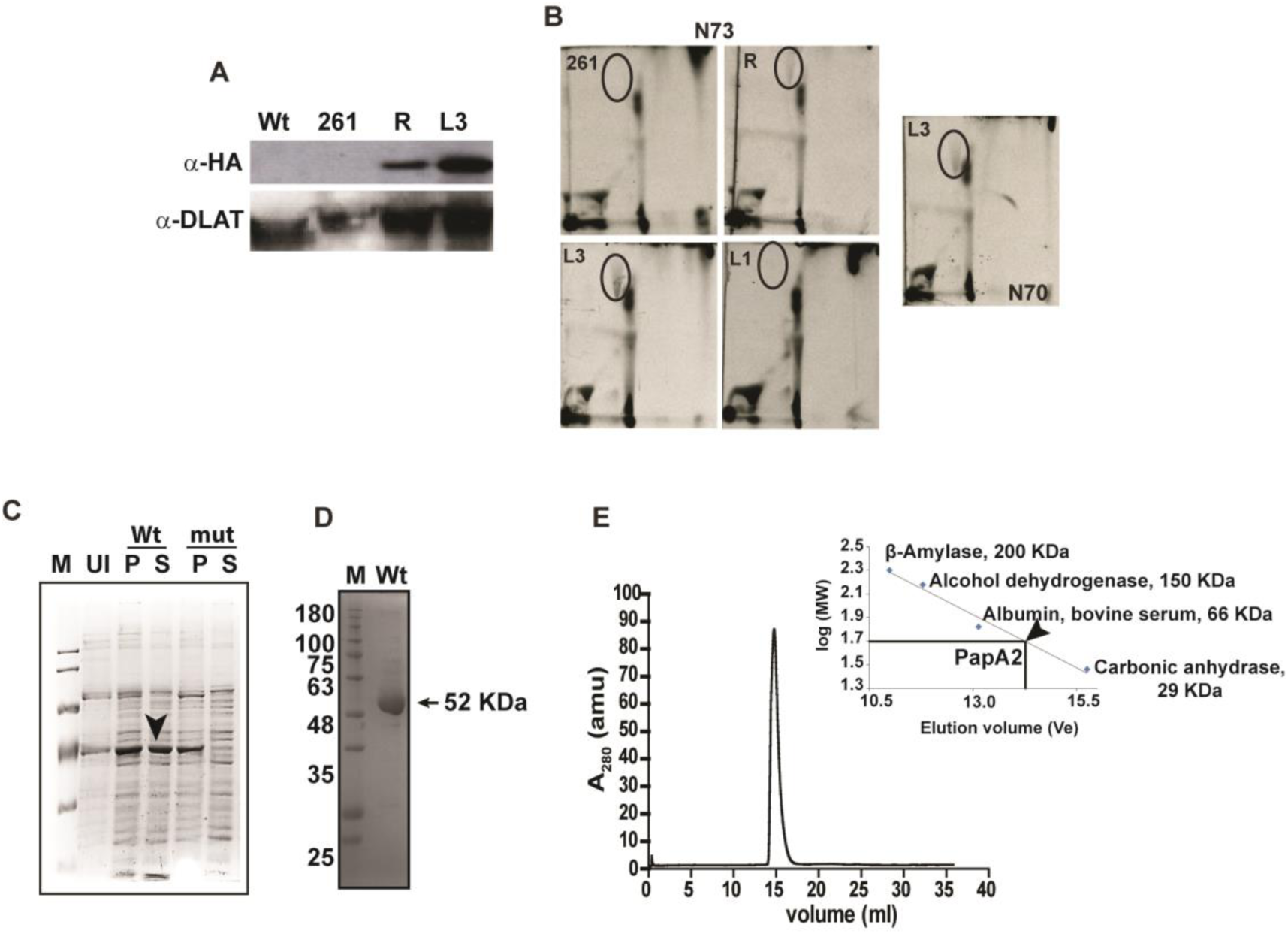
Functional complementation of SL-1 in L1 Mtb by PapA2 from the L3/ L4 Mtb strains. A) Analysis of PapA2 expression in extracts of N73 Mtb by immunoblotting with HA or DLAT antibodies. 261-denotes extracts from N73 transformed with pMV261 alone. R and L3 refer to N73 transformed with PapA2 from H_37_Rv and N24 strains, respectively. B) Analysis of apolar lipids from PapA2 expressing Indo oceanic L1 Mtb strains. B-Lipids from ^14^C acetate labelled cultures analysed in solvent D by TLC. The apolar lipids from another Indo oceanic L1 strain (N70) transformed with L3 PapA2 is also shown. The SL-1 spot is encircled. C) Expression profile of Wt-PapA2 and A83P-PapA2 was analysed at indicated time points post-induction using 100 µM IPTG through SDS-PAGE. Induction experiments were performed at 18°C. M is protein ladder, Wt and mut represents wild type and A83P –PapA2, U is uninduced protein sample, P and S are pellet and soluble fraction of induced protein samples. Black and white arrows indicate expression of Wt and mutant PapA2 in the soluble fraction, respectively. D) Analysis of the purified Wt PapA2 by SDS-PAGE. E) Analysis of oligomeric state of the purified Wt PapA2 by Gel exclusion chromatography. The elution profile of Wt papA2 is depicted; the elution volumes of standard proteins used for mass calculation are indicated in the inset graph.

To test if this mutation affected the structural integrity of PapA2 or its function, we expressed both the Wt and mutant proteins in *E. coli*. While we could obtain ∼ 50% soluble protein expression of the Wt, the mutant protein partitioned to insoluble fractions in all conditions of culture and expression (Fig. 2C) suggestive of a strong influence of the mutation on overall protein structure of PapA2. We resorted to a 2-step approach to confirm the role of this mutation in protein structure-1) establish structure of the Wt protein and 2) understand the effect of A83P substitution by molecular dynamics simulation. A significantly pure in excess of 90% of native PapA2 in the monomeric state (Fig. 2F, G) was subjected to X-ray crystallography for determination of structure after confirming the veracity of the purified protein by MALDI-TOF mass spectrometry (Fig. S1).

### PapA2 structural features display an unusual NRPS C domain architecture

The structure of PapA2 was determined at resolution of 2.16 Å using the phase calculated from anomalous diffraction of selenium as described in Methods (PDB ID-6AEF). Details of data collection and data processing are summarized in Table 2. The asymmetric unit possess two molecules (Fig. 3A). Each monomeric structure can be further described by dividing the protein into two subdomains: an N-terminal subdomain (residues 2-215) and a C-terminal subdomain (residues 216-459), each with a classical CAT fold comprising of a large β sheet flanked by alpha helices (Fig. 3B). The core β sheet in the N-terminal subdomain encompasses seven mixed type beta strands (parallel and antiparallel) - β1, β2, β3, β6, β7, β8 and β13 whereas the C-terminal subdomain contains six mixed beta strands -β9, β10, β11, β12, β14 and β15 (Fig. 3B). The two subdomains are connected by two cross-over points – “latches”, wherein the C-terminal subdomain extends back to the N-terminal subdomain - 1) “N-terminal latch” - residues 311-323, forming helix-α14 and 2) “C-terminal latch” -residues 391- 411, including helix- α18 followed by a beta strand- β13 (Fig. 3B). Most importantly, structure analyses indicated the presence of a unique Zn binding motif (ZnF) in the N-terminal subdomain comprising of residues-C19, H21, H124, H135 (Fig. 3C). The presence of Zn in the native protein was further confirmed through ICP-MS.

**Table 1.**
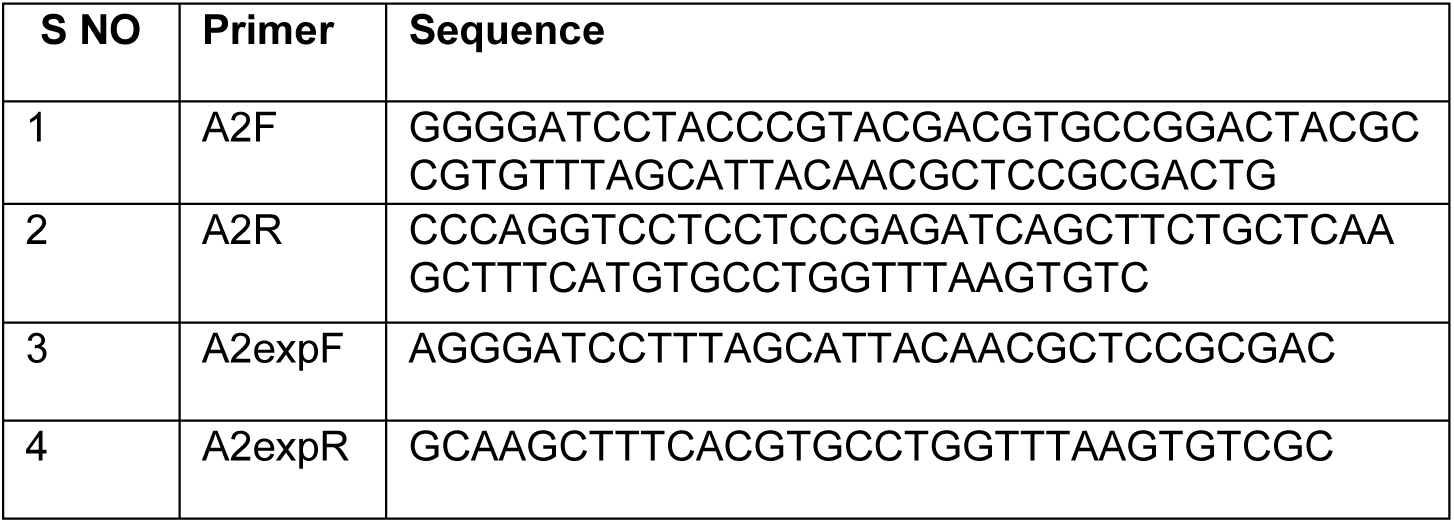
List of primers used in this study.

**Table 2.** Crystallographic data collection and refinement statistics.

| Dataset | Native | SeMet |
| --- | --- | --- |
| **Data collection** |  |  |
| Space group | P2<sub>1</sub>2<sub>1</sub>2<sub>1</sub> | P2<sub>1</sub>2<sub>1</sub>2<sub>1</sub> |
| Cell dimension |  |  |
| a, b, c (Å) | 74.71, 100.7, 128.9 | 74.67, 97.49, 128.8 |
| α, β, γ (°) | 90°, 90°, 90° | 90°, 90°, 90° |
| Resolution (Å) | 50- 2.16 (2.25- 2.16) | 50- 2.5 (2.54- 2.5) |
| Redundancy | 6.5 (3.5) | 8.7 (8.3) |
| R<sub>merge</sub> | 9.5 (81.5) | 11.7 (61.3) |
| CC (1/2) (%) | 78.7 | 90.4 |
| <I/σI> | 19.7 (2.24) | 22.09 (3.59) |
| Completeness (%) | 99.5 (96.2) | 96.2 (97.1) |
| **Refinement** |  |  |
| Resolution (Å) | 39.71-2.16 |  |
| No. of reflections | 52058 |  |
| R<sub>work</sub>/R<sub>free</sub> | 0.1981/0.2315 |  |
| No. of atoms | 14,660 |  |
| Protein | 14,159 |  |
| Water | 472 |  |
| Zn | 2 |  |
| ACT | 7 |  |
| TRS | 20 |  |
| B factors | 46.0 |  |
| Protein | 32.1 |  |
| Water | 36.8 |  |
| RMSD |  |  |
| Bond lengths (Å) | 0.005 |  |
| Bond angles (°) | 0.711 |  |

**Fig. 3.**
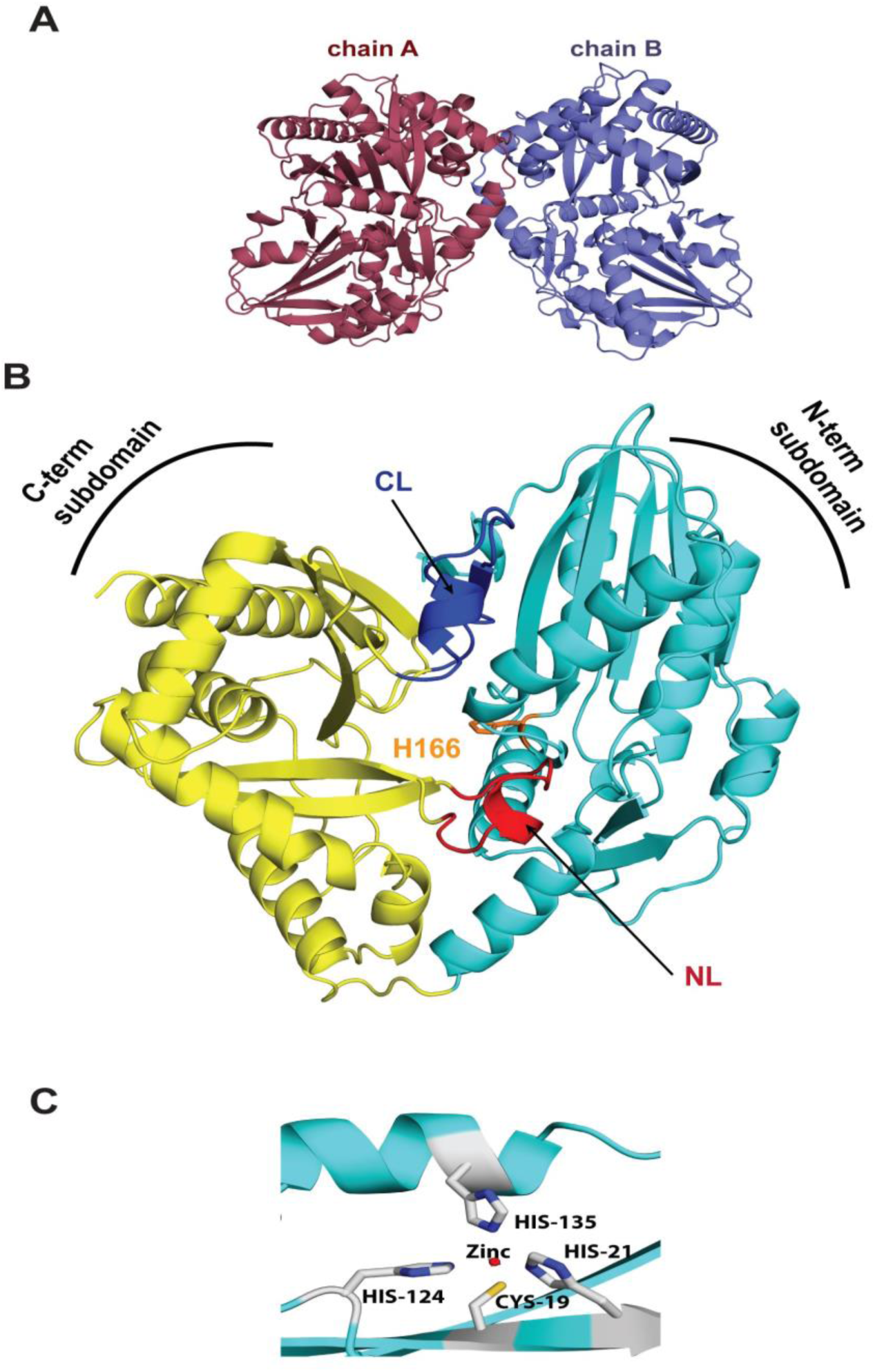
PapA2 portrays a two- domain architecture. A) Asymmetric unit of the crystal contains two molecules of PapA2-chain A and B. B) The overall structure of PapA2. N-terminal subdomain is cyan, C-terminal subdomain is golden yellow, N-terminal and C-terminal latches are shown in red (NL) and blue (CL) colour respectively. The location of active site H166 in orange colour. C) The putative Zn finger motif is shown-Zn atom (red circle) and coordinating residues (grey).

### The interface region of PapA2 possess substrate binding sites

Solid surface analysis revealed the presence of a solvent accessible tunnel at the interface of two subdomains (Fig. 4A). Placement of pseudo-tunnel at the interface of two subdomains using Caver program (30) and metapocket server (31) showed H166, the catalytic centre (32), in the middle of the tunnel indicating a potential substrate binding site(s). This ∼ 25 Å long tunnel originated close to the “N-terminal latch” and ended before helix 3 with an access to His166 from both ends will henceforth be referred to as the “tunnel” (Fig. 4B).

**Fig. 4.**
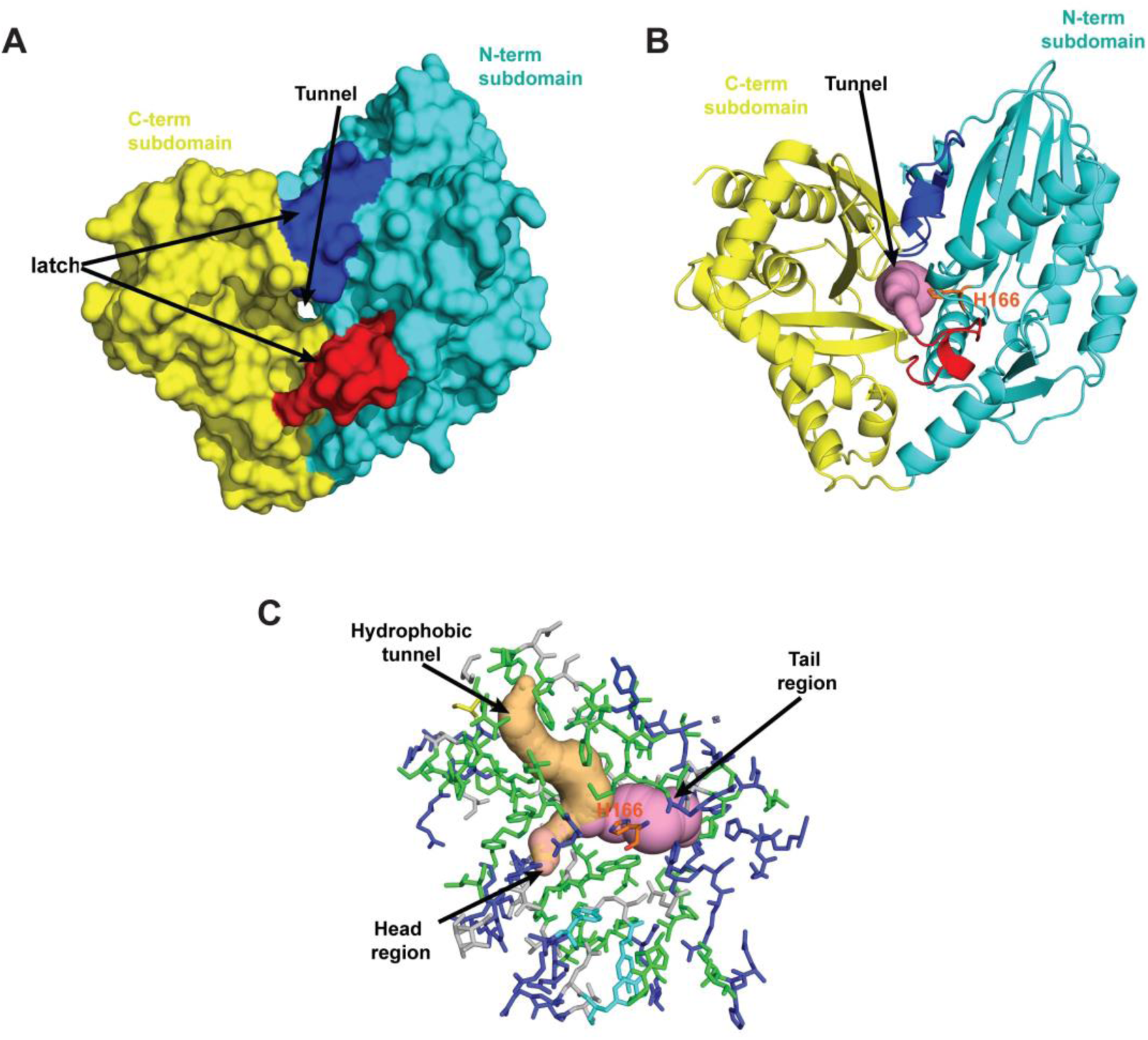
The Interface region of two subdomains harbours a tunnel for putative substrate binding. A, B) Solid surface (A) and cartoon presentation (B) of PapA2 protein structure. The tunnel (pink) is seen at the centre of interface regions of two subdomains-N-terminal subdomain (cyan) and C-terminal subdomain (golden yellow). The two crossover points or latches are shown red and blue. C) Stick presentation of the interface region along with pseudo tunnels displaying void regions in the protein. The tunnel (pink) and hydrophobic tunnel (light orange) are marked. Residues lining the interface region are colour coded based on their polarity; hydrophobic (green), charged (tv blue) and polar (grey). Active site is shown in orange.

Further detailed analysis revealed a well-organized arrangement of residues with distinct polarity at the “interface” tunnel with a dense population of positively charged arginine residues in the tail region of the tunnel and a more diverse distribution of residue polarity in the head region of the tunnel (Fig. 4C). Interestingly, another cavity enriched in hydrophobic residues at one end (the “hydrophobic tunnel”) intersects with the tunnel in close proximity to the catalytic H166 suggesting a putative binding site for the large hydrophobic acyl chain of the donor substrate.

### Molecular docking provides evidence for a unique substrate approach to PapA2

We further investigated the interface region by molecular docking using the acceptor and donor substrates. We reasoned that the catalytic H166 should be in close proximity of the acylation site of the substrate for efficient catalysis. Using this as a reference, we selected the conformer that positioned the 2’-OH of Trehalose-2-Sulfate in the proximity of H166 and also identified 4 residues-P171, T307, T324 and-S384 in apposition of the ligand (Fig. 5A). Similar docking studies also clearly placed the long acyl chain of palmitoyl CoA (donor substrate) in the hydrophobic tunnel and the CoA moiety in proximity to the tail region of the open tunnel (Fig. 5B) revealing a putative unique bidirectional substrate approach to the catalytic centre of PapA2 - access of the acceptor substrate from the head region of the open tunnel and entry of donor substrate from the tail region during the acylation reaction.

**Fig. 5.**
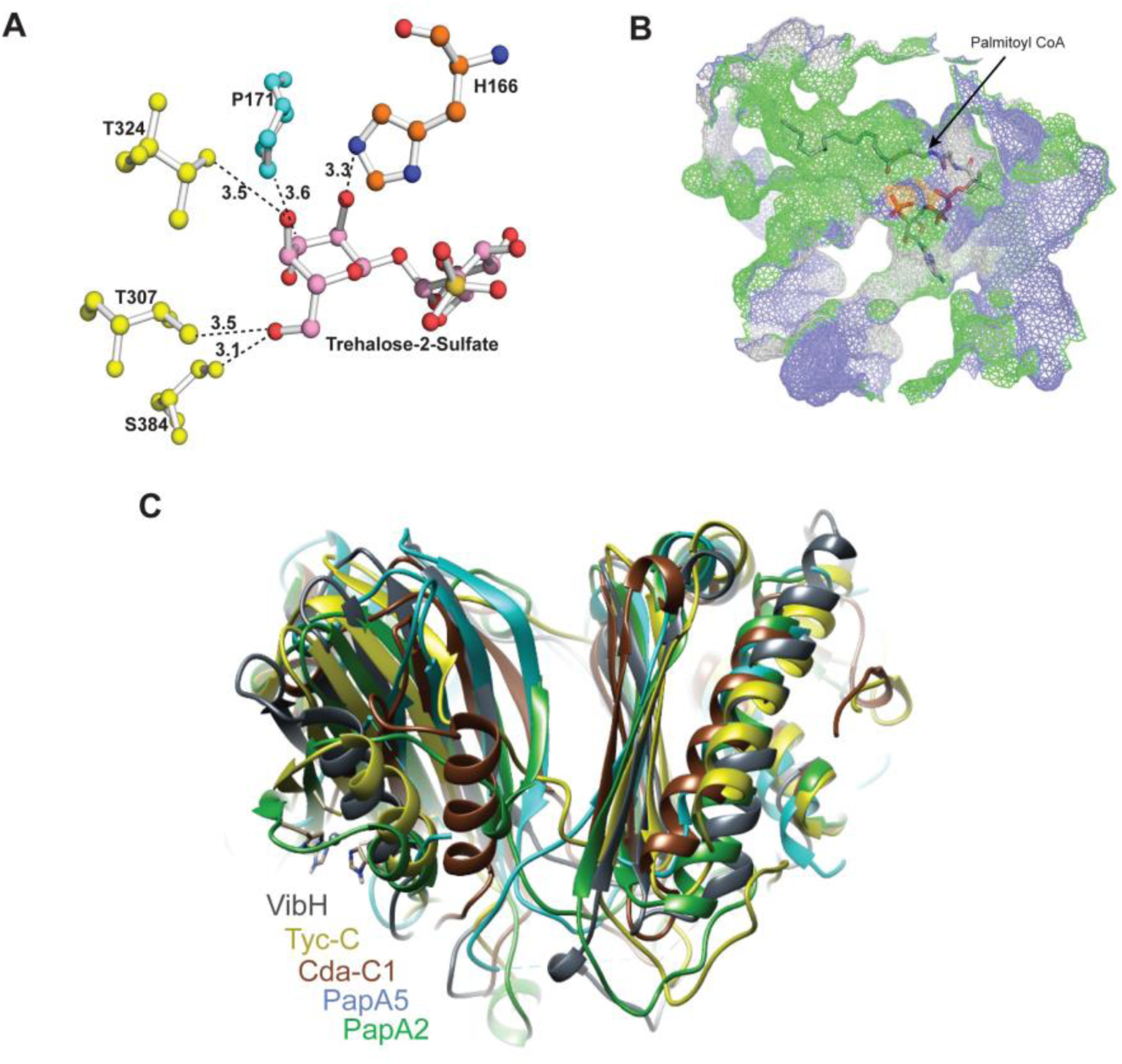
Detailed analysis of the acceptor and donor substrate binding sites. A) Ball and stick model of the substrate binding sites. The 2’-OH group of acceptor substrate is in proximity to ε nitrogen of H166 of condensation motif, with P171, T307, T324 and S384 being additional residues in proximity to the acceptor substrate. B) Mesh surface representation shows the donor substrate conformer with the acyl component residing in a hydrophobic tunnel and the CoA component in close proximity of the tail region of the solvent accessible tunnel. C) Superposition of PapA2 with other stand-alone C domain structures revealed a distinct NRPS C-domain architecture.

In an effort to define the substrate binding region, we superimposed PapA2 with previously reported structures of other C domain proteins-PapA5 (Polyketide associated protein of acyltransferase-5) (33), CDA-C1 (Condensation (C)-domain of -Calcium Dependent Antibiotic synthetase) (34), TycC (Tyrocidine synthetase III) (35), SrfC (Surfactin A synthetase C) (36) and VibH (Vibriobactin synthase) (37). Although, we observed an overall conservation of architecture of the proteins with the catalytic histidine residing in the subdomain interface region, and conserved positioning of secondary elements in the C terminal subdomain, considerable conformational differences were observed in the N terminal subdomain (Fig. 5C). Previous studies have identified the key acceptor substrate determinants for some of these proteins by mutagenesis (34, 37, 38). Our docking studies identified important residues of the head region of the solvent accessible tunnel - P171, T307, T324, S384 residing in close proximity of the acceptor substrate (Fig. 5A). Surprisingly, mapping with the other C-domain proteins revealed three of four residues of Mtb PapA2 (P171, T324 and S384) as positional equivalents of acceptor substrate determinants in VibH (G131 and N335) or CDA-C1 (G162, S309) or PapA5 (G129) (Table 3).

**Table 3.** Molecular determinants for acceptor substrate from various C domain proteins.

| VibH | Cda-C1 | PapA5 | PapA2 |
| --- | --- | --- | --- |
| G131 | G162 | G129 | P171 |
| - | S309 | - | T324 |
| N335 | - | - | S384 |
| H126 | H157 | H124 | H166 |
| D130 | D171 | D128 | D170 |

### MD simulations reveal putative local and global changes in A83P- mutant PapA2

MD Simulations have been extensively used in the past to understand intrinsic dynamics behaviour for various proteins. Together with crystallographic data, this complementary approach captures dynamics and structural insights of protein conformational state. To generate the starting structure of mutant, we modelled the A83P residue site of PapA2 structure (Fig. 6A). Interestingly, the mutation is distal to other functional sites, including ZnF motif and catalytic motif, and was mapped to the protein surface (α4 helix) (Fig. 6B). Two independent MD simulations of the wild-type and mutant PapA2 were performed for a cumulative 1 μs simulation time. Standard MD protocol, including all-atomistic representation was used (see Methods for details).

**Fig. 6.**
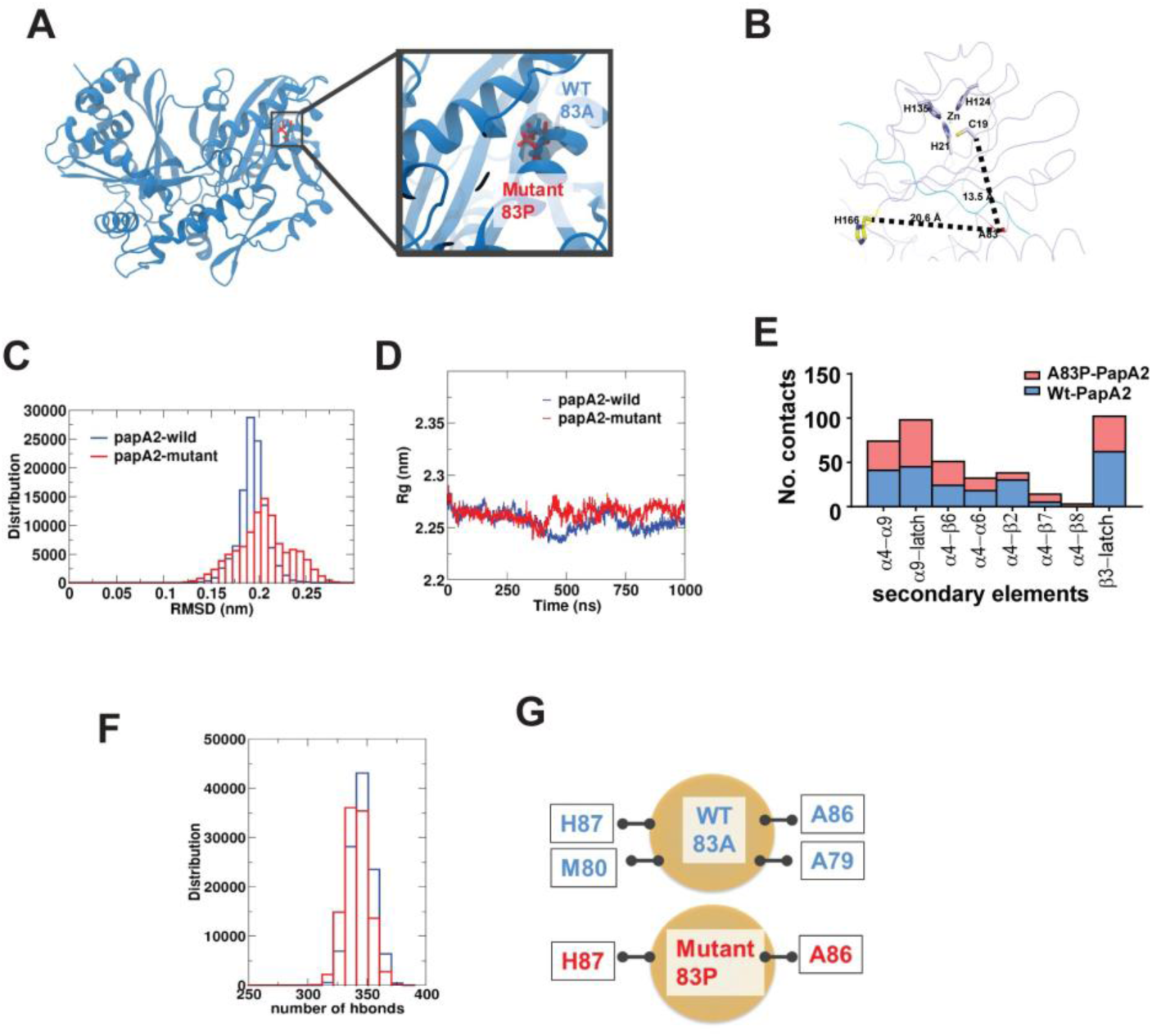
Molecular dynamic simulation reveals major alterations in the global fold of mutant PapA2. A) Representation of the models (Wt and Mutant PapA2) structures used for MD simulation studies. B) Distance of the A83P mutation from the H166 catalytic residue and the Zn finger motif of PapA2. C) Graphical representation of the RMSD changes in Wt and Mutant PapA2. D) Analysis of changes in the total size (radius of gyration –Rg) of PapA2 protein on account of the A83P mutation. E) Analysis of numbers of local contacts of the 83^rd^ residue in Wt (blue) and A83P (red)PapA2. The local secondary elements are represented on the x-axis. F, G) Analysis of total (F) hydrogen bond interactions in the Wt (blue) and A83P (red) and hydrogen bonds interacting with the 83^rd^ residue of Mtb PapA2.

Comparison of wild-type and mutant trajectories revealed a large number of global structural rearrangements (Fig. 6C-D). The order parameters, RMSD and Rg capture the mobility and overall compactness of the protein, respectively. The mutant simulations showed increased mobility with protein ensembles exhibiting >0.2 nm RMSD compared with the observed crystallographic wild-type structure. The resultant increase in Rg at ∼400ns pointed towards a change in protein conforrmation. Interestingly, a significant increase in the flexibility of the C-terminal latch (> 0.5 nm) in A83P-PapA2 was again supportive of the distal effects of the A83P mutation on PapA2 flexibility and overall the global fold of the protein.

Concomitant to global changes, we also monitored local dynamical changes induced by the mutant residue. Analysis of neighbouring residues that directly contact residue 83 of PapA2 identified major alterations. Local contacts were significantly reduced for the α4- α6, α4- α2 and β3- latch regions with increase in contacts the α4- β7 region as a result of the proline mutation in mutant PapA2 (Fig. 6E). In addition, while A83 bonded with H87, M80, A86, and A79 via hydrogen bonds, analysis of the P83 interactions significantly reduced the local network to specifically two bonds (H87, and A86) (Fig. 6F, G), lending further support to the substantial local destabilization of non covalent interactions that serves as nucleation site to unfold the protein (the global change) and affect overall protein stability.

## Discussion

The mycobacterial cell wall is one of the most complex chemical entities replete in lipid and carbohydrate moieties not found elsewhere in biological systems apart from actinomycetes. The intricate molecular mechanisms for the biosynthesis, maintenance and plasticity is yet not well understood; more so the adaptability in the face of physiological stress and environmental pressures. Given the long evolution of Mtb, its adaptation with the human host in the context of its genomic and molecular repertoire is now being recognised as an important factor for the successful survival. Recent molecular evidences have pointed out lineage specific variations in Mtb, culminating as a result of both host driven and environmental cues, correlating with the inflammatory potential of the pathogen (1, 39, 40). Often, a clear correlation between the observed lineage specific genetic attributes and its molecular / phenotypic characteristics or vice versa remains poorly characterized.

In an effort to understand the molecular basis for loss of sulfolipid in clinical strains of Mtb (a small subset of Mtb lineage 1), we identified a single SNP in the deficient strains mapping to the PapA2 coding region of the genome. This resultant conversion of the 83^rd^ alanine to proline of PapA2, one of the primary enzymes of the mycobacterial sulfolipid biosynthetic machinery manifested as a compromise in protein stability and folding, significant enough to prevent our attempts to obtain purified, soluble mutant protein even by chemical chaperones. This global defect in the fold of the mutant protein was further supported by MD simulation studies and emphasizes a crucial role for the A83 residue in structural integrity of PapA2.

In agreement with the structure of other C domain multidomain peptide synthases proteins of NRPS (non-ribosomal peptide synthases) (33, 34, 36, 37), PapA2 also conformed to a typical V shaped two subdomain containing architecture with two latch components and catalytic histidine at the interface of two subdomains. The distinct presence of a solvent accessible tunnel in close apposition with the catalytic site and a hydrophobic tunnel implies a dual substrate approach strategy- an acceptor substrate (T2S) approaching from the head region and the donor acyl CoA substrate from the tail region (acyl group residing in the hydrophobic rich region) and the CoA in close proximity of the tail region. Remarkably, the A83P mutation mapped to the protein surface away from the catalytic and Zinc finger containing regions (20 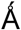 and 13 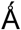 respectively) of L1 PapA2. Our conclusion of mutation site distant from the active site deteriorates protein structure and function is in agreement with the reports on the effect of distant mutation in the proteins (41, 42). Interestingly, the occurrence of hydrophobic residues, such as Ala or Val, in this position is highly conserved in all the members of the PapA proteins of Mtb suggestive of the importance of this Ala residue in the function of this protein family (Fig. 7).

**Fig. 7.**
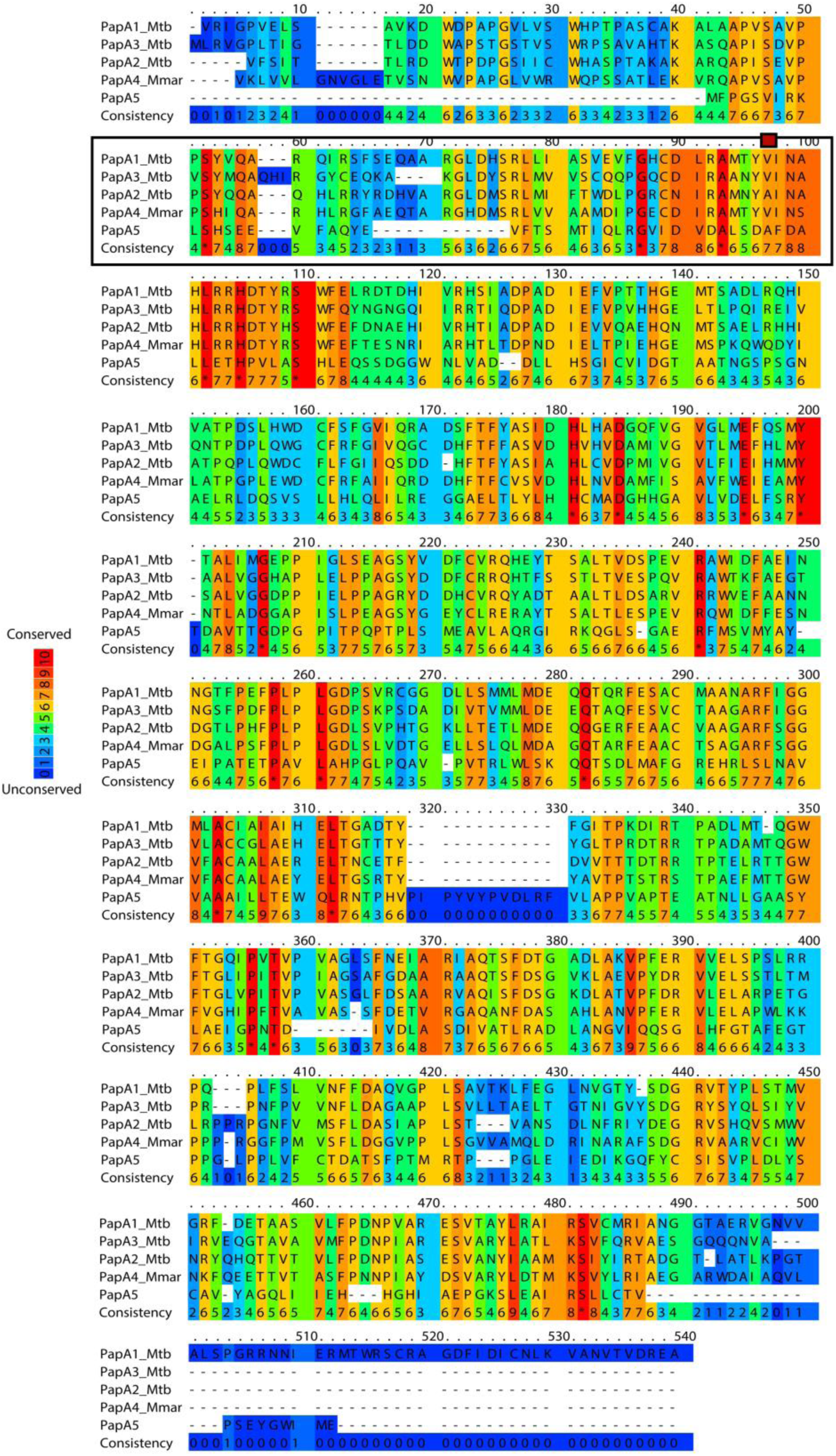
Multi-sequence alignment places the PapA2 83^rd^ residue in a highly conserved region of the Mtb PapA protein family. The protein sequence of PapA1-5 proteins of Mtb were aligned by Clustal W (degree of conservation is indicated as a colour scale). The conserved region encompassing the 83^rd^ residue is outlined by a box.

The most interesting aspect of the structure was the identification of a putative Zn finger motif in PapA2. Presence of this motif implores the possibility of protein-protein/ protein – DNA interactions unique to any of the acyl transferase proteins identified till date. While PapA2 is involved in the first acylation of T2S, PapA1 catalyses the addition of a second acyl chain to this mono-acylated sulphated trehalose. We identified a long hydrophobic tunnel that could house the acyl CoA donor necessary for PapA2 function. However, PapA1 function requires 2 acyl chains to be fitted in close proximity of each other (acyl group of mono-acylated – T2S and the second fatty acyl CoA). It is logical to assume the possibility of an interaction between PapA2 and PapA1 in order to facilitate this dual acyl chain transfer. The presence of the Zn finger motif in native PapA2 strongly suggests a similar protein-protein interaction in mycobacterial sulfolipid biosynthesis. In fact, previous studies have proposed a similar multi component- “scaffolding” model for the biosynthesis of SL-1, which require biosynthetic components in close proximity to each other (3). Similar mechanisms of protein-protein mediated transfer of long chain acyl moieties has been reported for the synthesis of another complex glycolipid-TDM in mycobacteria wherein antigen 85 complex proteins mediate the inter protein transfer of mycolate chains from two trehalose mono-mycolates to form one molecule of trehalose dimycolate (27, 29, 43). Identifying the interactions between the PapA proteins of mycobacteria would provide conclusive evidence of such novel functions associated with cell wall lipid assembly in Mtb.

The role of sulfolipids in Mtb pathogenesis is confusing. Classical studies have linked the expression of sulfolipids to the degree of virulence associated with Mtb (27). However, Rousseau et.al, 2003 have clearly unlinked the presence of sulfolipid with virulence in H_37_Rv (44). While the importance of sulfolipid in host- pathogen cross talk can be envisaged given its localization to outer most layer of the cell, it has also been implicated in inhibition of phago-lysosome fusion (21) and modulating the pro-inflammatory response (22, 23). Alternatively, by functioning as a sink to buffer changes in propionyl CoA content SL can contribute to metabolic reshuffling during *in-vivo* growth (16, 45, 46). In contrast, mutants of H_37_Rv that lack sulfolipids viz. Δpks2, Δmmpl8 have not shown any defect in *in vivo* growth in mice models of infection (7, 47). Given the pleomorphic importance of PGL in Mtb virulence and its dependence on the strain genotype, it is plausible to expect that this specific loss of sulfolipid −1 in the Indo-oceanic L1 strains specifically is an adaptive mechanism for the fine-tuned balance of infection by these strains in the specific human population/ environment. A careful elucidation of the importance of SL-1 in mycobacterial immuno-pathogenesis in the context of specific lineages would aid resolve this conundrum. Our phenotype-genotype correlation of a novel SNP resulting in the loss of a major surface glycolipid in a specific subset of Mtb provides an excellent platform to address specific adaptive mechanisms employed by a very successful human pathogen.

## MATERIAL AND METHODS

### Bacterial cell cultures

*Mycobacterium tuberculosis* strains were grown in Middlebrook 7H9 (BD Biosciences, USA) media containing ADC (Albumin Dextrose Catalase) (BD Biosciences, USA) at 37°C under shaking conditions unless stated otherwise. The Mtb clinical strains (Table S1) were a kind gift of Dr. Sebastien Gagneux, Swiss TPH and part of the San Francisco collection (1). *E. coli* strains were cultured as per standard procedures in LB broth or agar (BD Biosciences, USA). Media was supplemented with kanamycin (50μg /ml) or carbenicillin (100μg/ml) when needed.

### Analysis of lipids from Mtb

10 ml Mtb grown to the logarithmic phase in 7H9 media was supplemented with 1 µCi of ^14^C-acetate (American Radiolabeled Chemicals, Inc. USA) for 24h at 37°C following which the polar and apolar lipids were isolated according to standard protocols (43). The extent of radiolabel incorporation was estimated by using TopCount NXT scintillation counter (PerkinElmer, Ohio, USA). Lipids equivalent to 10000 cpm for all the three strains were spotted on TLC silica gel 60 (Merck Millipore, USA) and eluted using solvents (A-D) for apolar lipids and (D, E) for polar lipids (43). The TLCs were developed onto X rays or scanned using GE Typhoon FLA 7000 phosphorimager system (GE Healthcare Bio-Sciences, USA).

### Cloning, Expression and Purification of Mtb-PapA2

For mycobacterial expression of PapA2, the complete ORF of *papA*2 was PCR amplified from the genomic DNA of H37Rv (R), N24 (L3) or N73 (L1) and cloned into the mycobacterial expression vector pMV261 by using specific primers-A2F and A2R to express the recombinant protein as a HA tagged fusion protein. For expression in *E. coli*, *papA*2 was amplified from the genomic DNA of H37Rv (Wt PapA2) or N73 (mutant PapA2) using primers A2expF and A2expR, cloned into pET28-SMT3 vector to obtain the plasmid pVIP06. A list of primers is given in Table 1. Expression of recombinant protein as tested in the *E. coli* strain C41(DE3) with isopropyl-β-D-thiogalactoside (IPTG) (Himedia laboratories, India). For seleno-methionine labelled protein (SeMet-PapA2), cultures were grown at 25°C in selenoMet Dream Nutrient Mix (Molecular Dimensions, UK). Large scale purified protein was obtained from cultures induced with 0.1mM IPTG for 24 h at 18°C by using affinity columns (Ni-NTA agarose, Qiagen, Germany). The protein was eluted with 250 mM of Imidazole (Himedia laboratories, India), concentrated using Amicon Ultra Centrifugal Filters (Merck life sciences, Germany) and subjected to Ulp1 protease at 4°C for 16h for tag removal. Further purifications Superdex-75 10/300gl (GE Healthcare Life Sciences, UK) and anion exchange chromatography using a Resource Q column (GE Healthcare Life Sciences, UK) resulted in a > 90 % pure protein. Expression of PapA2 was confirmed by immunoblotting with Tag specific antibody-(ab18181-HA/ ab18184-His, Abcam, UK).

### Sample preparation and MALDI-TOF mass spectrometry

For MALDI-TOF, 25µg of PapA2 protein subjected to trypsin digestion was injected into the MALDI-TOF mass spectrometer-in a MALDI-TOF/TOF 5800 (AB Sciex, USA) and the fragments were identified from SwissProt.

### Protein crystallization

Sparse matrix crystallization trials of PapA2 (at 10 mg/ml) were carried out with Crystal Screen HT (Hampton research, CA, USA) by Hanging drop vapour diffusion technique (48) at 25°C. Initial diffraction experiments were performed using crystals obtained in 0.2 M MgCl_2_·4H_2_O, 0.1 M C_2_H_12_AsNaO_5_·3H_2_O, pH 6.5, 20 % w/v PEG 8,000 buffer. Following further optimization, the crystals were stored frozen with with 30 % w/v PEG 8,000 as a cryoprotectant.

### Data collection and processing

Diffraction data for native PapA2 and SeMet-PapA2 crystals were collected at European Synchrotron Radiation Facility (ESRF, Grenoble, France) on the beam line BM14. PapA2 and SeMet-PapA2 crystals were diffracted up to 2.16 Å and 2.49 Å resolutions, respectively. Data obtained were Indexed and scaled using the program HKL-2000 (49). The scaled intensities were converted to structure factors using the program TRUNCATE (DOI-10.1107/S0567739478001114) as implemented in CCP4 (50). The phase problem was solved using the selenium as heavy atoms and by applying Single Anomalous Dispersion (SAD) phasing procedure using AutoSol wizard in PHENIX (51, 52). The structure of PapA2, was determined by Molecular replacement (53) using chain (B) of SeM-PapA2 as template in Phaser (54) in PHENIX, refined as rigid body followed by restraint refinement using phenix.refine (55). The model was built into the electron density map using the program COOT (56). The program PyMOL (PyMOL Molecular Graphics System, Schrödinger, LLC) was used to visualize and analyse the model.

### Molecular Dynamic Simulation

The crystal structure of papA2 was cleaned and prepared using Maestro (Schrödinger) (Maestro, version 9.8, Schrödinger, LLC, New York, NY, 2014). The prepared structure was selected for generation of mutation, A83P, using Accelrys Discovery Studio visualizer (Discovery Studio. “version 2.5.” Accelrys Inc.: San Diego, CA, USA (2009). Both the Wt and mutant PapA2 structures were taken up for further refinement by molecular dynamics (MD) simulations by GROMACS and OPLS-all atom force field (DOI-10.1021/ja9621760) using Wt papA2 as the starting structure. Both structures were used for separate MD simulations for 1μs each. Each of the starting structure was placed in a cubic box solvated using TIP4P water representation (DOI-10.1080/00268978500103111) (see Table 4). The systems were neutralized using Na+ ions. The starting structures were subjected to energy minimization using the steepest descent method. Systems were simulated at 300K using Nose-Hoover T-coupling (DOI-10.1063/1.447334) and then later subjected to Parrinello-Rahman barostat (DOI-10.1063/1.328693) for pressure coupling at 1 bar, before the production run were started. Electrostatic interactions were calculated using the particle mesh Ewald (PME) summation (DOI-10.1063/1.464397).

**Table 4:**
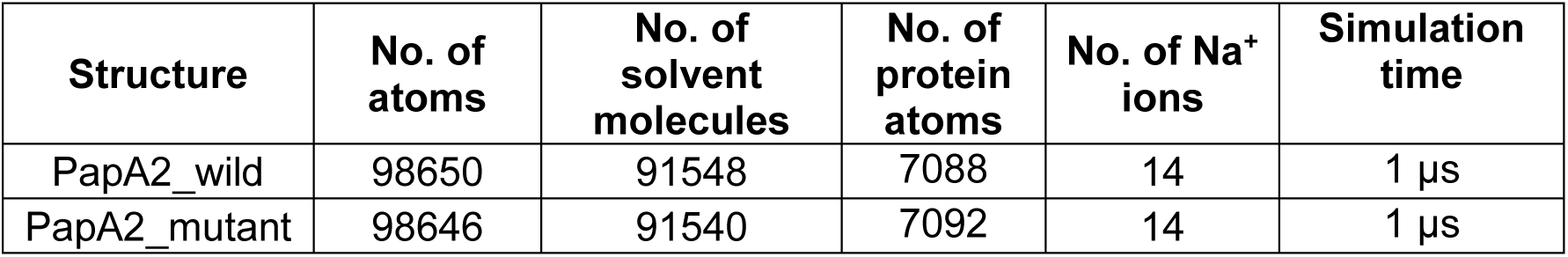
Details of MD simulation of Wt and mutant PapA2.

| Structure | No. of atoms | No. of solvent molecules | No. of protein atoms | No. of Na<sup>+</sup> ions | Simulation time |
| --- | --- | --- | --- | --- | --- |
| PapA2_wild | 98650 | 91548 | 7088 | 14 | 1 μs |
| PapA2_mutant | 98646 | 91540 | 7092 | 14 | 1 μs |

## Acknowledgments

The authors acknowledge National Institute of Immunology and Regional Centre for Biotechnology for in house beamline facility and ESRF beamline BM14 for diffraction data collection. The Mosquito robotic system, mass spectrometry central facilities at CSIR-IGIB are duly acknowledged. VR acknowledges grants from CSIR (BSC0123, BSC0124), SC acknowledges the DST SERB Ramanujan fellowship SB/S2/RJN-14/2013. VP acknowledges UGC-JRF for Ph.D. Fellowship. LT and NJ acknowledge CSIR-4PI for supercomputing facilities. LT acknowledges the support from DST-INSPIRE Faculty funded research grant [LSBM-45] from Department of Science and Technology, India. NJ is thankful to DBT-RA and DST-NPDF Fellowships. Proof reading by Prabhakar A and Deepthi S is duly acknowledged.

## Author Contributions

VR conceptualized the work, VP, VR, SC, AB, GB were involved in the design of the work. VR and AB were involved in the mycobacterial work. VP and AM were involved in protein purification and crystallization, VP determined and analysed the structure, NJ performed the molecular simulations, VP and NJ analysed the molecular simulation results. VP, LT, VR and SC have contributed to the manuscript preparation.

## Conflict of interest

The authors do not have any competing interests.

---

**Fig. S1.**
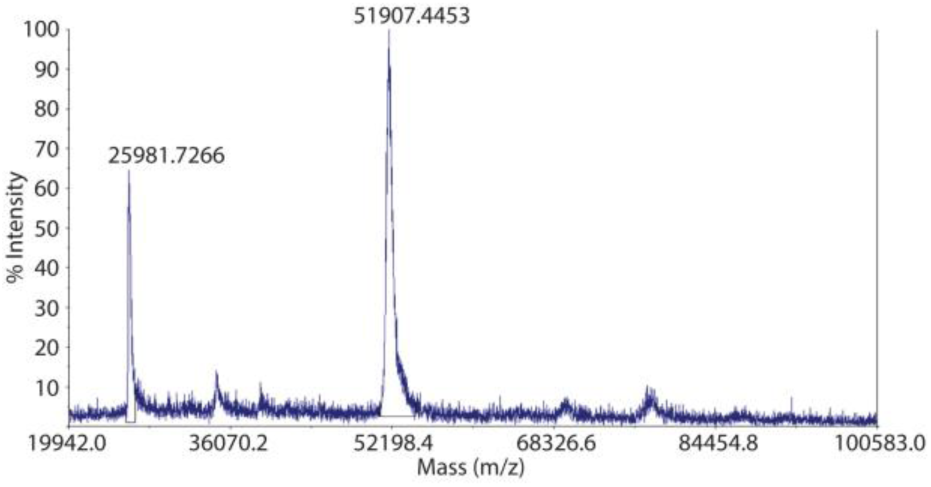
Confirmation of the identity of purified PapA2 by MALDI-TOF. The MALDI-TOF spectra of purified PapA2 is shown. The peak at the expected molecular mass of 52kda confirms the identity of the protein.

**Table S1.** Summary of Mtb strains used in this study.

| Strain name | Lineage | Origin |
| --- | --- | --- |
| N70 | 1 | San Francisco, USA |
| N72 | 1 | San Francisco, USA |
| N73 | 1 | San Francisco, USA |
| T83 | 1 | San Francisco, USA |
| N04 | 3 | San Francisco, USA |
| N24 | 3 | San Francisco, USA |
| N37 | 3 | San Francisco, USA |
| Erdman | 4 | Jeff Cox, USA |

